# COFFEE: Consensus Single Cell-Type Specific Inference for Gene Regulatory Networks

**DOI:** 10.1101/2024.01.05.574445

**Authors:** Musaddiq K Lodi, Anna Chernikov, Preetam Ghosh

## Abstract

The inference of gene regulatory networks (GRNs) is crucial to understanding the regulatory mechanisms that govern biological processes. GRNs may be represented as edges in a graph, and hence have been inferred computationally for scRNA-seq data. A wisdom of crowds approach to integrate edges from several GRNs to create one composite GRN has demonstrated improved performance when compared to individual algorithm implementations on bulk RNA-seq and microarray data. In an effort to extend this approach to scRNA-seq data, we present COFFEE (COnsensus single cell-type speciFic inFerence for gEnE regulatory networks), a Borda voting based consensus algorithm that integrates information from 10 established GRN inference methods. We conclude that COFFEE has improved performance across synthetic, curated and experimental datasets when compared to baseline methods. Additionally, we show that a modified version of COFFEE can be leveraged to improve performance on newer cell-type specific GRN inference methods. Overall, our results demonstrate that consensus based methods with pertinent modifications continue to be valuable for GRN inference at the single cell level.

## 1 Introduction

The study of biological systems is being conducted in several different ways. One popular way of analyzing the relationship between chromatin, transcription factors and genes is to represent them as a complex network known as the gene regulatory network (GRN). GRNs are crucial to the understanding of how cellular identity is established and corrupted in disease. The popular abstraction for analyzing GRNs is in the form of graphs, in which the relationship between any two genes is quantified by an edge score. The goal of GRN inference is to better understand the gene expression patterns that connect transcription factors and signaling proteins to target genes [1]. Several algorithms have been developed to infer GRNs from bulk RNA-sequencing data [2][3][4]. However, single cell transcription data presents the opportunity to observe cell-type specific gene expression patterns and potentially gain further insight into the regulation of cells [5]. The noise present in scRNA-seq datasets makes it difficult to determine if results are biologically significant. Hence, validation of constructed GRNs through pathway enrichment and literature review become paramount [6].

Algorithms that infer GRNs from bulk RNA-sequencing data have been adapted for single cell transcriptome data, to varying degrees of success. The algorithms available to the community vary in architecture and approach. Some are correlation based methods, such as LEAP and PPCOR [7] [8]. More recent algorithms rely on linear regression and non-linear ordinary differential equations to make GRN predictions. Additionally, some algorithms require the input of time-point expression data, while others do not. Often times, scRNA-seq experiments do not collect this information, so a common accepted practice has been to generate pseudotime point data with methods such as Slingshot [9]. The increasing number of algorithms for this purpose becomes an insurmountable task for researchers; how should a scientist choose the best algorithm to construct a GRN from single cell transcriptomics data? As scRNA-seq data has become more accessible, Pratapa et al. sought to solve this problem by creating an evaluation framework for twelve prominent GRN inference algorithms [2]. Based on the performance of these algorithms on synthetic, curated and experimental single cell transcriptomic datasets, the authors were able to make the recommendation that the algorithms PIDC, GENIE3, and GRNBoost2 are the methods of choice for researchers seeking to use a GRN inference algorithm [2]. The robustness of these algorithms was measured using the early precision ratio (EPR) score. EPR is the early precision ratio of a network, which essentially measures the number of true positive interactions in a network [10]. Rather than choosing select algorithms based on an evaluation criteria, another approach to constructing GRNs is to leverage information from all of the present algorithms, using the wisdom of crowds theory.

The wisdom of crowds theory states the collective knowledge of a community is greater than the knowledge of any individual. This theory has broad practical applications to a number of fields [11]. Its implementation in GRN inference is not a new concept; a study in 2012 by the DREAM5 Consortium et al. used a consensus network approach for bulk RNA-sequencing data [12]. This consensus approach used the Borda count method, which is a ranked choice voting algorithm invented by John Charles de Borda in 1770. The way the system works is that candidates are ranked by the choice of the crowd, and are assigned a score accordingly (the candidate ranked in first receives the maximum number of points, and so on). The candidate with the highest average score then wins the election [13]. Building on this platform, we implemented a normalized version of the Borda count system to generate a consensus network approach applied to the inference of miRNA-miRNA networks [14] [15]. This current study seeks to improve this implementation by making it more specific to GRN inference for single cell transcriptomics data.

The consensus approach for GRN inference has demonstrated improved performance for microarray and bulk RNA-seq data [12]. Our motivation for this study was two-fold: we wanted to test if a similar wisdom of crowds approach is effective for scRNA-seq data. GRN inference for scRNA-seq data presents a unique set of challenges. The noise present in pseudotime data, as well as gene/cell dropouts, requires more sensitive algorithms to predict high quality interactions [16]. With sub-optimal performance shown in several GRN inference algorithms, integrating them to achieve increased performance may not necessarily be as intuitive as inference for bulk-RNA sequencing data [2]. Additionally, with the advent of higher quality scRNA-seq and scATAC-seq data, inferring GRNs by individual cell-type has become a paramount task. Determining the individual regulatory interactions per cell-type can lead to the identification of novel cell-types and disease progression mechanisms [17]. We sought to determine if a consensus-based approach can lead to higher performance in GRN inference for specific cell-types. To this end, we present COFFEE, a consensus algorithm for scRNA-seq data, for both cell-type specific and general scRNA-seq data. Since the consensus approach works on the individual networks predicted by each algorithm, it can readily integrate algorithms that leverage scRNA-seq data or even multi-omics datasets (e.g., scRNA-seq and scATAC-seq data).

## 2 Methods

### 2.1 Selection of Algorithms

The algorithms selected for this study that form the basis for the consensus network construction were from the BEELINE framework proposed by Pratapa et al. A summary of these algorithms is presented in Table 1.

**Table 1:**
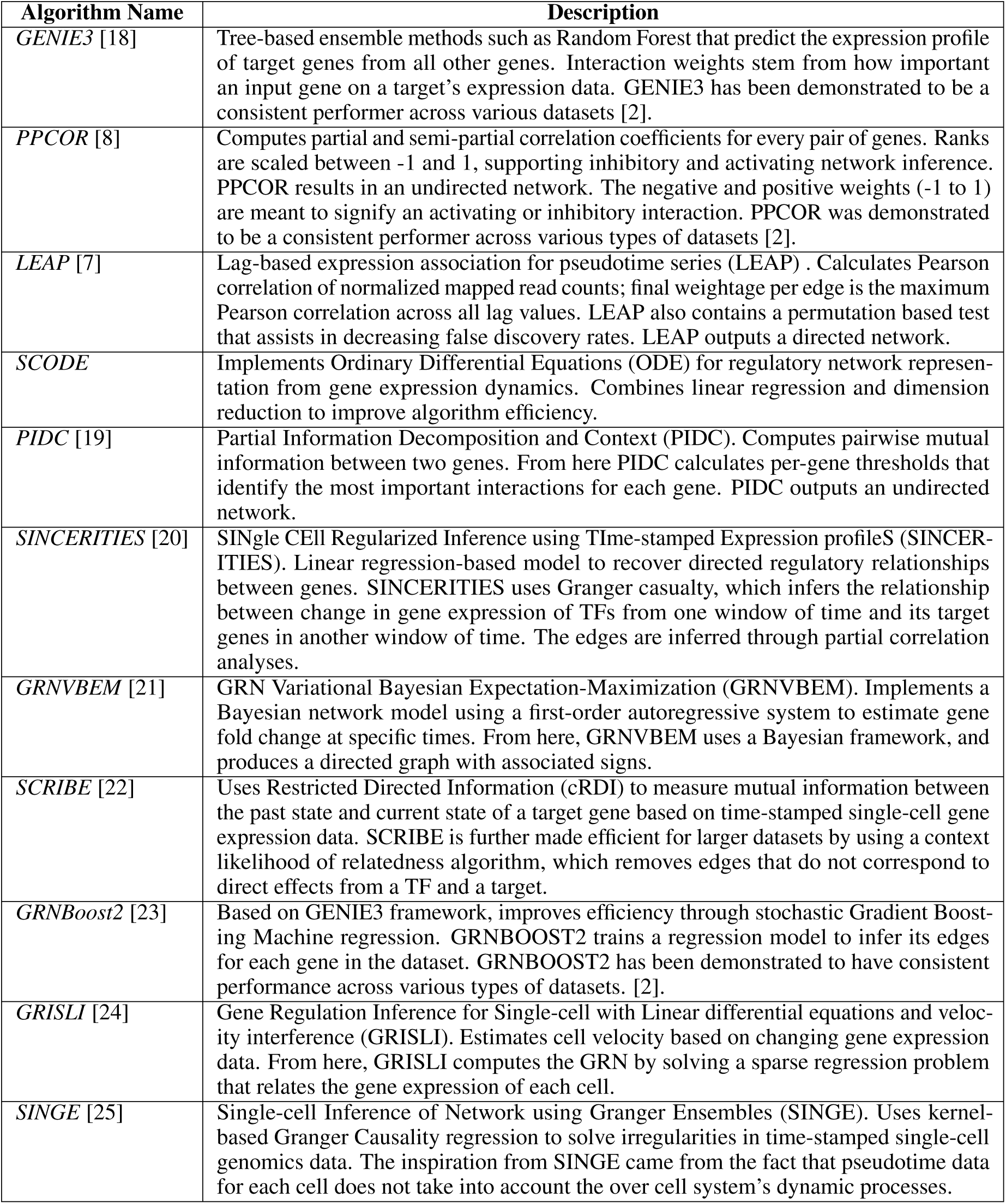
Summary of algorithms used in the BEELINE framework that were leveraged in our proposed consensus network approach.

| Algorithm Name | Description |
| --- | --- |
| *GENIE3* [18] | Tree-based ensemble methods such as Random Forest that predict the expression profile of target genes from all other genes. Interaction weights stem from how important an input gene on a target’s expression data. GENIE3 has been demonstrated to be a consistent performer across various datasets [2]. |
| *PPCOR* [8] | Computes partial and semi-partial correlation coefficients for every pair of genes. Ranks are scaled between -1 and 1, supporting inhibitory and activating network inference. PPCOR results in an undirected network. The negative and positive weights (-1 to 1) are meant to signify an activating or inhibitory interaction. PPCOR was demonstrated to be a consistent performer across various types of datasets [2]. |
| *LEAP* [7] | Lag-based expression association for pseudotime series (LEAP) . Calculates Pearson correlation of normalized mapped read counts; final weightage per edge is the maximum Pearson correlation across all lag values. LEAP also contains a permutation based test that assists in decreasing false discovery rates. LEAP outputs a directed network. |
| *SCODE* | Implements Ordinary Differential Equations (ODE) for regulatory network representation from gene expression dynamics. Combines linear regression and dimension reduction to improve algorithm efficiency. |
| *PIDC* [19] | Partial Information Decomposition and Context (PIDC). Computes pairwise mutual information between two genes. From here PIDC calculates per-gene thresholds that identify the most important interactions for each gene. PIDC outputs an undirected network. |
| *SINCERITIES* [20] | SINGLE CELL Regularized Inference using Time-stamped Expression profileS (SINCERITIES). Linear regression-based model to recover directed regulatory relationships between genes. SINCERITIES uses Granger casualty, which infers the relationship between change in gene expression of TFs from one window of time and its target genes in another window of time. The edges are inferred through partial correlation analyses. |
| *GRNVBEM* [21] | GRN Variational Bayesian Expectation-Maximization (GRNVBEM). Implements a Bayesian network model using a first-order autoregressive system to estimate gene fold change at specific times. From here, GRNVBEM uses a Bayesian framework, and produces a directed graph with associated signs. |
| *SCRIBE* [22] | Uses Restricted Directed Information (cRDI) to measure mutual information between the past state and current state of a target gene based on time-stamped single-cell gene expression data. SCRIBE is further made efficient for larger datasets by using a context likelihood of relatedness algorithm, which removes edges that do not correspond to direct effects from a TF and a target. |
| *GRNBoost2* [23] | Based on GENIE3 framework, improves efficiency through stochastic Gradient Boosting Machine regression. GRNBOOST2 trains a regression model to infer its edges for each gene in the dataset. GRNBOOST2 has been demonstrated to have consistent performance across various types of datasets. [2]. |
| *GRISLI* [24] | Gene Regulation Inference for Single-cell with Linear differential equations and velocity interference (GRISLI). Estimates cell velocity based on changing gene expression data. From here, GRISLI computes the GRN by solving a sparse regression problem that relates the gene expression of each cell. |
| *SINGE* [25] | Single-cell Inference of Network using Granger Ensembles (SINGE). Uses kernel-based Granger Causality regression to solve irregularities in time-stamped single-cell genomics data. The inspiration from SINGE came from the fact that pseudotime data for each cell does not take into account the over cell system’s dynamic processes. |

We implemented the BEELINE evaluation framework and ran each algorithm through the pipeline provided [2]. Once these networks for each individual algorithm were constructed, we used our proposed COFFEE framework to integrate them into one consensus network.

### 2.2 Borda Count Implementation

Each algorithm from the BEELINE implementation outputs a ranked edge list with a confidence score attached to each edge. Since each algorithm is normalized in a different way, the edge weights are not distributed equally, so the ranking of each edge within the list across algorithms is considered. For example, consider the ranking of four edges for three different algorithms, each inferring a distinct ranked edge list.

The Borda count method allocates points to each rank, where the highest ranked interaction receives the maximum number of points, and the lowest ranked interaction receives zero points. To receive a final rank between 0 and 1, the resulting weighted ranks are normalized. In the example in Tables 2 and 3, there are four example interactions; the interaction I4 is ranked at the first position for Algorithm 1. Therefore it receives the maximum of three Borda points, and a normalized score of 1. The resulting Borda rank used is the normalized number of points received for each algorithm.

**Table 2:** Ranked individual predictions for three example algorithms for example interactions denoted I.

| Normalized Borda Points (Borda Ranking) | Borda Points | Rank | Alg. 1 | Alg. 2 | Alg. 3 |
| --- | --- | --- | --- | --- | --- |
| 1 | 3 | 1 | *I4* | *I2* | *I2* |
| 0.667 | 2 | 2 | *I2* | *I3* | *I3* |
| 0.334 | 1 | 3 | *I1* | *I4* | *I1* |
| 0 | 0 | 4 | *I3* | *I1* | *I4* |

**Table 3:** Final Borda ranks for each example interaction.

| Interaction | Average of Borda Ranking | Final Rank |
| --- | --- | --- |
| *I2* | $(0.667+1+1) / 3$ | 0.889 |
| *I3* | $(0+0.667+0.667) / 3$ | 0.445 |
| *I4* | $(1+0.334+0) / 3$ | 0.445 |
| *I1* | $(0.334+0+0.334) / 3$ | 0.222 |

**Table 4:** Experimental Datasets Used to Evaluate COFFEE.

| Dataset | Description |
| --- | --- |
| hHEP [32] | scRNA-seq experiment on induced pluripotent stem cells (iPSC). Contains 425 scRNA-seq measurements across various timepoints. Pseudotime was calculated using Slingshot with day 0 as the starting cluster and day 21 as the ending cluster [9]. |
| hESCs [33] | Timecourse scRNA-seq experiment from 758 cells using the differentiation protocol in order to produce definitive endoderm cells from human embryonic stem cells, measured at 0, 12, 24, 36, 72 and 96 hours (h). Using slingshot, the pseudotime was calculated using 0h as the starting cluster and 96h as the ending cluster [9]. |
| mHSCs [34] | Contains normalized expression data for 1,656 HSPCs across 4,773 genes. Pseudotime was computed using Slingshot across three lineages, which were erythroid, granulocyte-monocyte and lymphoid [9]. The GRN for each lineage was separately inferred. |
| Mouse embryonic stem cells (mESC) [35] | scRNA-seq expression measurements for 421 primitive endoderm (PrE) cells differentiated from mESCs, from five different time points: 0, 12, 24, 48, and 72 hours (h). Pseudotime was calculated using Slingshot, with the starting cluster being 0h and the ending cluster being 72h [9]. |

**Table 5:** Number of Cells per Cell Type in Human Fetal Hematopoiesis Cell Dataset.

| Cell-Type | Number of Cells |
| --- | --- |
| GPs-Granulocytes | 443 |
| HSC-MPP | 1367 |
| LMP | 1522 |
| MEMP | 1522 |

Modifications were made to the original Borda count method in order to make it more applicable for gene-gene relationships. Self loops in the graph were removed, so a gene interacting with itself was removed from consideration in the final consensus algorithm. Additionally, due to high variance amongst algorithms, there were several cases where an edge would appear in some algorithms with a high ranking, and not appear at all in others. To handle this, 0 was inserted where an edge was not present into the calculation and was considered for the Final Rank. We implemented a version of the Borda count algorithm in Python, which takes a directory of ranked lists as inputs, and will output the consensus network for a user specified threshold.

### 2.3 Single Cell-type specific GRN inference using multimodal data: scMTNI

A promising direction of GRN inference for single cell transcriptomics data is identifying unique GRN’s by cell type. The primary challenge of this approach is identifying accurate cell lineages, and the varying transcription factors that define them [17]. One way to infer high quality cell lineages, and therefore the important transcription factors by cell type, is to integrate scRNA-seq data with scATAC-seq data [5]. Single Cell Multi-Task Inference (scMTNI) is a recently published multi-task learning framework that integrates scATAC-seq with scRNA-seq data to infer cell-type specific GRNs. It has demonstrated improved performance over existing cell-type specific inference models. [5].

scMTNI is a multi-task learning framework that uses a probabilistic graphical model-based method to infer gene regulatory network dynamics from a cell lineage tree. The method defines a cell type as a group of cells with similar transcriptome and accessibility levels. Each cell type is treated as a task, and the goal of the method is to infer a GRN for each task, as well as the ideal parameters. scMTNI calculates the probability of each gene and its set of regulators for each cell type, and uses this probability calculation to inform the predicted network. For our comparative analysis, we used the inferred consensus networks for each cell type in the coarse Human Fetal Hematopoiesis dataset, with four cell types [26]. scMTNI was executed on several subsamples of each cell type, and the final network prediction was informed through a confidence score from the subsampling. We used a confidence cut off of 0.8 to calculate the evaluation metrics, as recommended by the authors of scMTNI [5].

### 2.4 Cell-Type Specific COFFEE Framework

The algorithms used for COFFEE have not been specifically optimized for cell-type GRN inference. However, interactions that are predicted from these algorithms may still be valuable in predicting cell-type specific GRNs. To test this theory, we used an adapted version of COFFEE and compared its performance to scMTNI on a cell-type specific dataset.

For cell-type specific GRN inference, we included four algorithms that do not require pseudotime data as input: PPCOR, GENIE3, GRNBOOST2, and PIDC [8] [18] [23] [19]. The reasons for this were two-fold. Firstly, these algorithms were shown to be the top performing and had high stability on experimental datasets [2]. Additionally, since they do not require pseudotime data as input, these algorithms were less likely to be sensitive to poor pseudotime calculation, and removes an element of uncertainty from the calculation.

When using COFFEE for cell-type specific GRN inference, we prioritized the information from the well established cell-type specific GRN inference algorithm, such as scMTNI. scMTNI is superior in inferring cell-type specific GRNs, as it incorporates scRNA-seq and scATAC-seq data [5]. We added scMTNI to the COFFEE pipeline, and calculated a consensus network with scMTNI, PPCOR, GENIE3, GRNBOOST2, and PIDC. For this analysis, we modified COFFEE to provide an initial score of 1 for any edges found in the scMTNI pipeline, thus prioritizing the edges found in scMTNI higher than that in the other four algorithms. Then, we evaluated COFFEE+scMTNI on the baseline inferred scMTNI network. Conceptually, this framework is similar to earlier consensus methods for GRN inference where expert knowledge on possible edges are prioritized a priori.

### 2.5 Filtering highly varying genes with Slingshot

The primary goal of Slingshot is to reconstruct pseudotime data based on cell lineages for scRNA-seq datasets. Slingshot organizes the cell into clusters and defines cell lineages based on the potential ordering or changing of cell states. We chose Slingshot to determine the pseudotime data to maintain consistency with the BEELINE evaluation framework; Pratapa et al. calculated the pseudotime data for the Synthetic, Curated and Experimental datasets used in our benchmarking, and we did not deviate from this calculation.

We used Slingshot to filter highly varying genes for cell-type specific GRN inference. We first loaded the gene expression matrix into the BEELINE package and performed the standard pre-processing and dimensionality reduction recommended by the package authors. Then, we calculated the highly varying genes across pseudotime points for one cell lineage, as we were computing varying genes per cell cluster. From here, we selected the top 500 highly varying genes; all pvalues were < 0.01 [9].

### 2.6 Evaluation

To evaluate our COFFEE framework against baseline algorithms, we used precision, recall, and F-score when comparing to the gold standard network.

#### 2.6.1 Precision

Precision is defined as the number of correctly predicted interactions divided by the total number of predicted interactions. It measures how accurate the positive predictions of the algorithm are and calculated as follows.

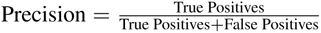

#### 2.6.2 Recall

Recall is defined as the ratio of all correctly predicted interactions to all actual positives. It measures how completely the algorithm predicts true interactions as follows.

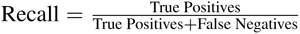

#### 2.6.3 F-Score

F-Score is defined as the harmonic mean between precision and recall. It provides a balanced representation of the relationship between precision and recall and calculated as follows.

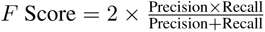

We used F-Score to determine the ideal threshold for COFFEE when applied to various dataset sizes.

## 3 Data

### 3.1 Synthetic Datasets

The primary benefit of using Synthetic datasets is to have a known GRN that has a comparable ground truth for reliable evaluation. Previous methods have used GeneNetWeaver; however, Pratapa et al. described limitations to this method [27]. Therefore, they created BoolODE. BoolODE converts a Boolean model to nonlinear ordinary differential equations which assists in capturing the logical relationship between regulators [2]. We directly used the Synthetic datasets provided by the authors of BEELINE without any additional preprocessing. The BEELINE authors created six datasets based on different network structures: Linear (LI), Cycle (CY), LL (Linear Long), BF (Bifurcating), BFC (Bifurcating Converging), and TF (Trifurcating). For our evaluation of COFFEE, we aggregated the datasets by number of cell-types, rather than the network structure. The cell-group sizes were 100 cells, 200 cells, 500 cells, 2000 cells and 5000 cells. Each group contained 50 individual datasets with a varying number of genes, each with their own ground-truth network. We used the Synthetic datasets to determine the ideal threshold for the consensus algorithm implemented in COFFEE.

### 3.2 Curated Datasets

Curated datasets are published Boolean models for GRNs that capture the specific regulatory processes of a given developmental process. For our evaluation, we selected three of the four Curated datasets used in the BEELINE evaluation framework: ventral spinal cord (VSC) development, hematopoietic stem cell (HSC) differentiation, and gonadal sex determination (GSD) [28] [29] [30]. The authors of the BEELINE framework used BoolODE to create 10 simulated datasets with 2000 cells based on the cell trajectories and gene expression patterns of the original Boolean models. We used the gene expression matrix and pseudotime data as is from the BEELINE data download, without any additional pre-processing.

The BEELINE authors also used the mammalian cortical area development (mCAD) Boolean model dataset in their evaluation framework; however, the authors noted that the algorithm results for this model were outliers, with poor performance results across all algorithms, which differed for the other Curated datasets [31] [2]. Therefore, we opted not to include mCAD in our evaluation process.

### 3.3 Experimental Datasets

To evaluate COFFEE on experimental methods, we chose two from human cells and two from mice cells. Similar to the Synthetic and Curated datasets, we used the pre-processed data from the BEELINE framework [2].

To compute the GRN inference using the important genes, we used the gene ordering file computed by the GAM R package to select the 500 top genes varying across pseudotime points, as detailed in the BEELINE evaluation protocol. This gene ordering file was provided in the dataset download from BEELINE. [2].

We also utilized the ground truth networks provided by the authors of BEELINE per experimental dataset cell type [2]. These ground truth sets were obtained from the ENCODE, ChIP-Atlas and ESCAPE databases for ChIP–seq data from the same or similar cell type. For our evaluation, we only considered interactions that contained genes present in the ground truth networks.

### 3.4 Cell Type Specific Dataset

To evaluate COFFEE’s performance on cell-type specific lineages, we used a dataset that scMTNI was benchmarked on. We compared the performance of COFFEE, scMTNI and COFFEE+scMTNI with a published scRNA-seq and scATAC-seq dataset of human fetal hematopoiesis cells. This study captured specifications for various blood lineages. We considered the coarse resolution of study to test the model on larger datasets, which contained four cell-types: g hematopoietic stem cell (HSC), multipotent lymphoid-myeloid progenitors (LMPs), MK-erythroid-mast progenitors (MEMPs), and granulocytic progenitors (GPs). The authors of scMTNI provided their pre-processed data per cell-type, in addition to the networks inferred by their method.

To use the data on COFFEE, we followed a similar process to our Experimental dataset evaluation by selecting the top 500 genes across pseudotime points using Slingshot [9]. To compare to scMTNI, we used the networks inferred by scMTNI provided in the author’s dataset download, using a confidence cutoff of 0.8, which is the recommendation of the authors of scMTNI [5].

## 4 Results

To evaluate the performance of the consensus algorithm using the Borda algorithm, we tested it on four different kinds of datasets: Synthetic, Curated, Experimental, and Cell Type Specific inference. In each case, we demonstrate that the wisdom of crowds approach leads to better performance across datasets. To evaluate, we used precision, recall, and F-score.

### 4.1 Synthetic Datasets

The Synthetic datasets were obtained from the Beeline evaluation framework [2]. We grouped the datasets by size, to evaluate the performance of the consensus algorithm as more genes and cells are present in a given dataset. There were five size groups present in the Synthetic datasets: 100 cells, 200 cells, 500 cells, 2000 cells, and 5000 cells. The number of genes varied depending on a specific dataset within the size group.

A key component of the consensus algorithm is determining a threshold at which to keep high confidence edges. A similar Borda based method for miRNA networks, miRsig used a default threshold cut off of 90%, which results in keeping the top 10% of predicted edges [14]. However, due to the cell to cell gene variation in gene expression present in single cell genomics data, we tested the algorithm on lower thresholds and evaluated its performance [36]. We used the mean F-Score to determine the ideal threshold value by dataset size. [14].

In Figure 1, we see that a different consensus threshold is appropriate depending on the size of the dataset. For the smaller datasets (100 or 200 cells), a threshold value of 0.75 leads to the best F-Score performance for COFFEE. For larger datasets (500, 2000 and 5000 cells), a threshold value of 0.65 leads to the best F-Score. Users may decide to maximize precision or recall rather than F-Score, which would lead to a different threshold being used. We found that increasing the threshold value increases the precision.

**Figure 1:**
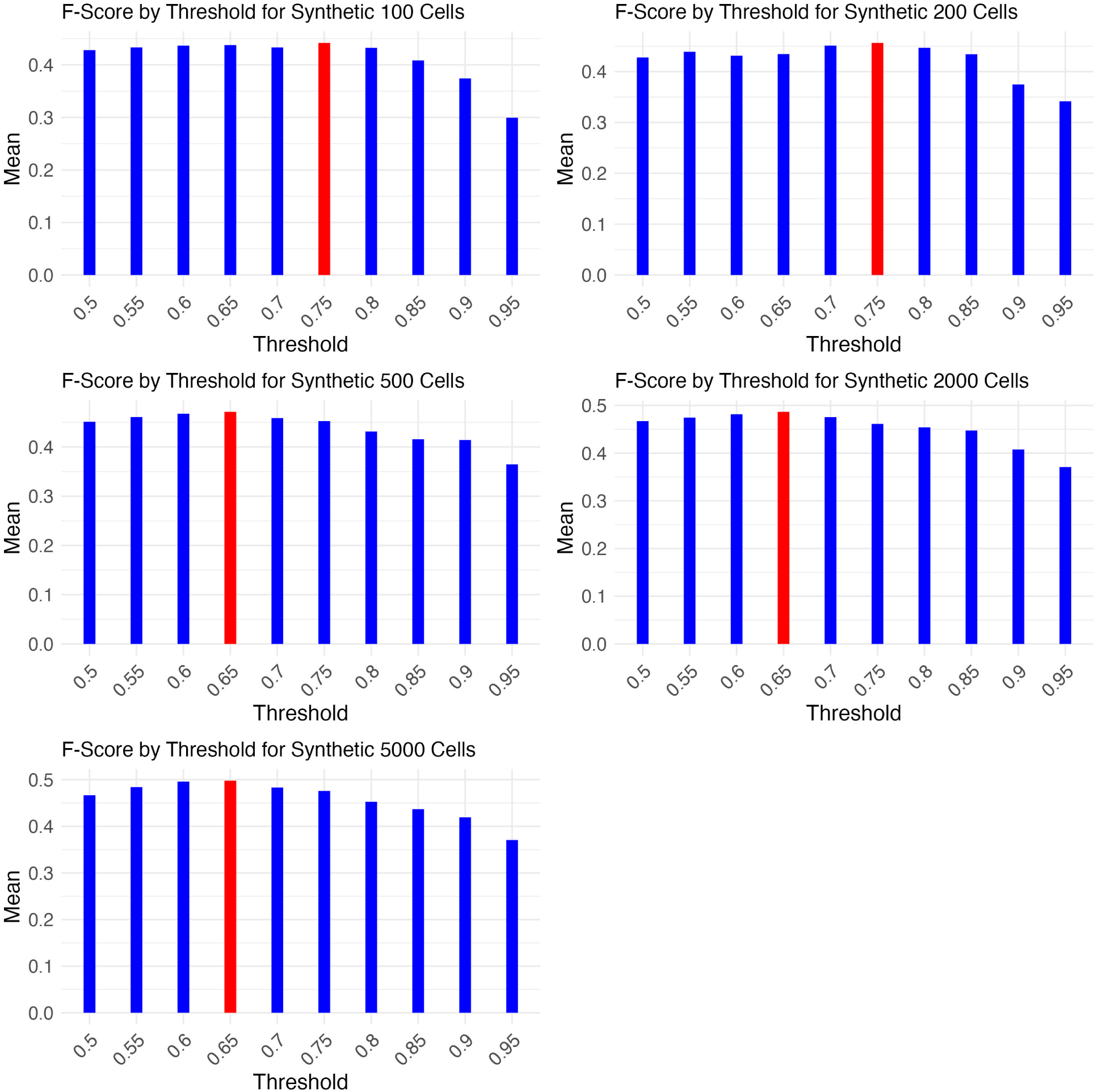
Optimal threshold for COFFEE based on dataset size, by mean F-Score. Highest mean F-Score is colored in red.

Table 6 shows the optimal threshold by F-Score for each dataset size. These thresholds were used for the evaluation of COFFEE against the baseline algorithms, as well as reporting the performance on Curated, Experimental and Cell-Type Specific datasets.

**Table 6:**
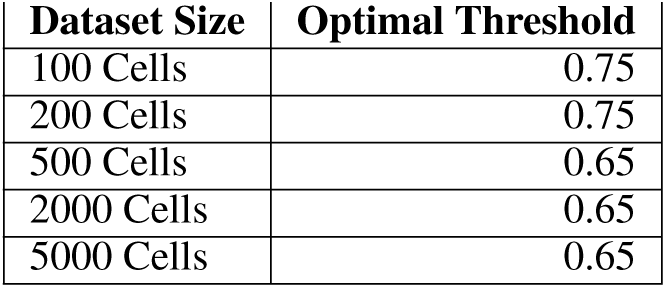
Optimal Threshold by F-Score for Synthetic Dataset Sizes.

| Dataset Size | Optimal Threshold |
| --- | --- |
| 100 Cells | 0.75 |
| 200 Cells | 0.75 |
| 500 Cells | 0.65 |
| 2000 Cells | 0.65 |
| 5000 Cells | 0.65 |

To evaluate the performance of COFFEE against the baseline algorithms, we primarily used F-Score, precision and recall as the metrics.

Figure 2 depicts the performance of COFFEE against the baseline algorithms by F-Score. We observe that across dataset sizes, COFFEE demonstrates a better performance. It is also significant to note that algorithms perform differently based on the data sizes. For example, SINCERITIES has a comparitively weaker F-Score for smaller datasets containing 100 or 200 cells than it does with 500, 2000 or 5000 cells. A consensus based approach such as COFFEE mitigates this variance by integrating information from the top performing algorithms despite dataset size. In short, COFFEE is less susceptible to variation in its performance based on differences present in a dataset.

**Figure 2:**
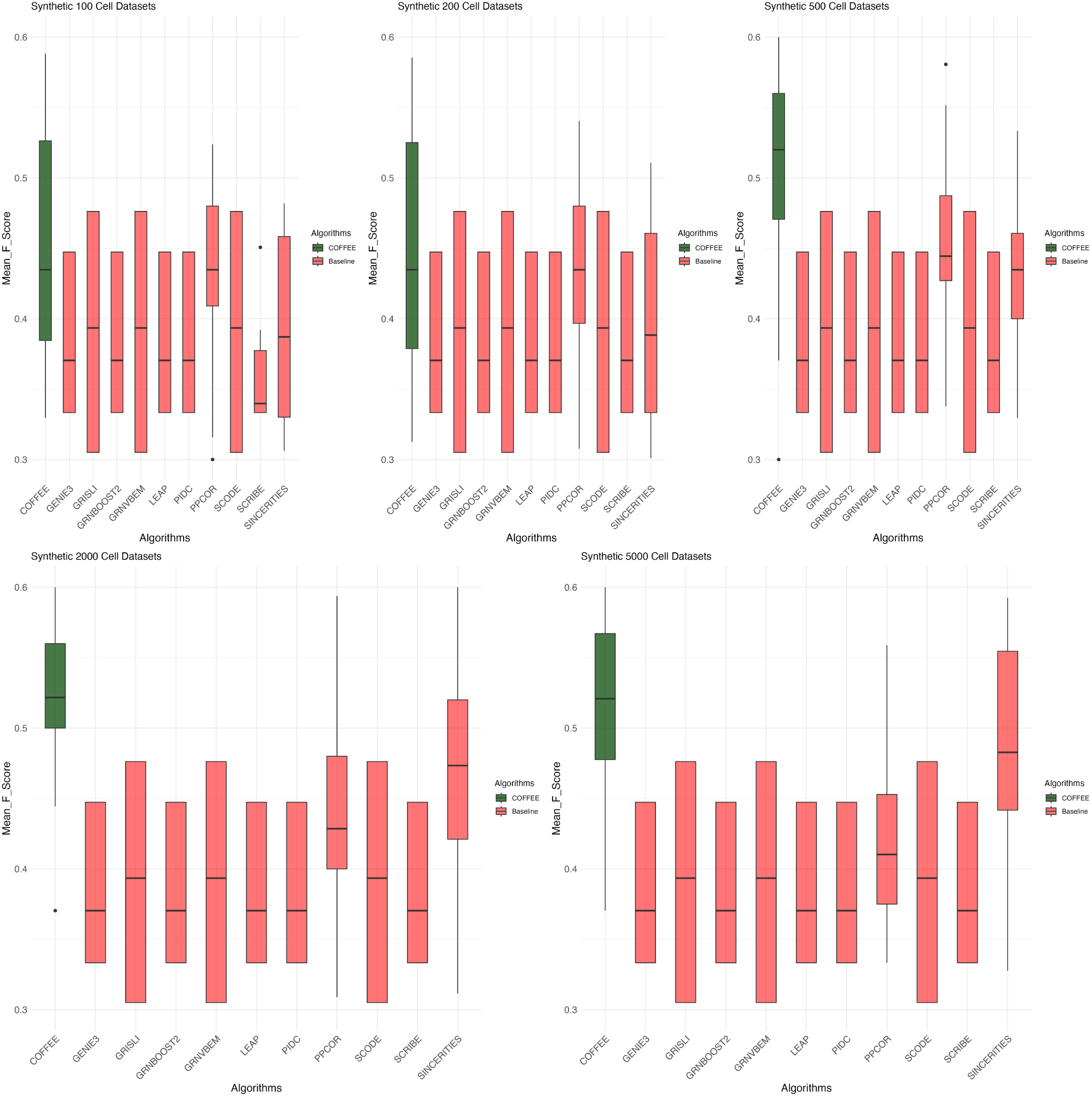
F-Score performance on Synthetic datasets grouped by size. COFFEE is colored in green.

To analyze the performance of the algorithms in further detail, we also looked at the mean precision and recall for each dataset size group across algorithms. From the analysis in Figure 3, we see that the precision in COFFEE is much higher than in any of the other base line algorithms, while its recall is much lower.

**Figure 3:**
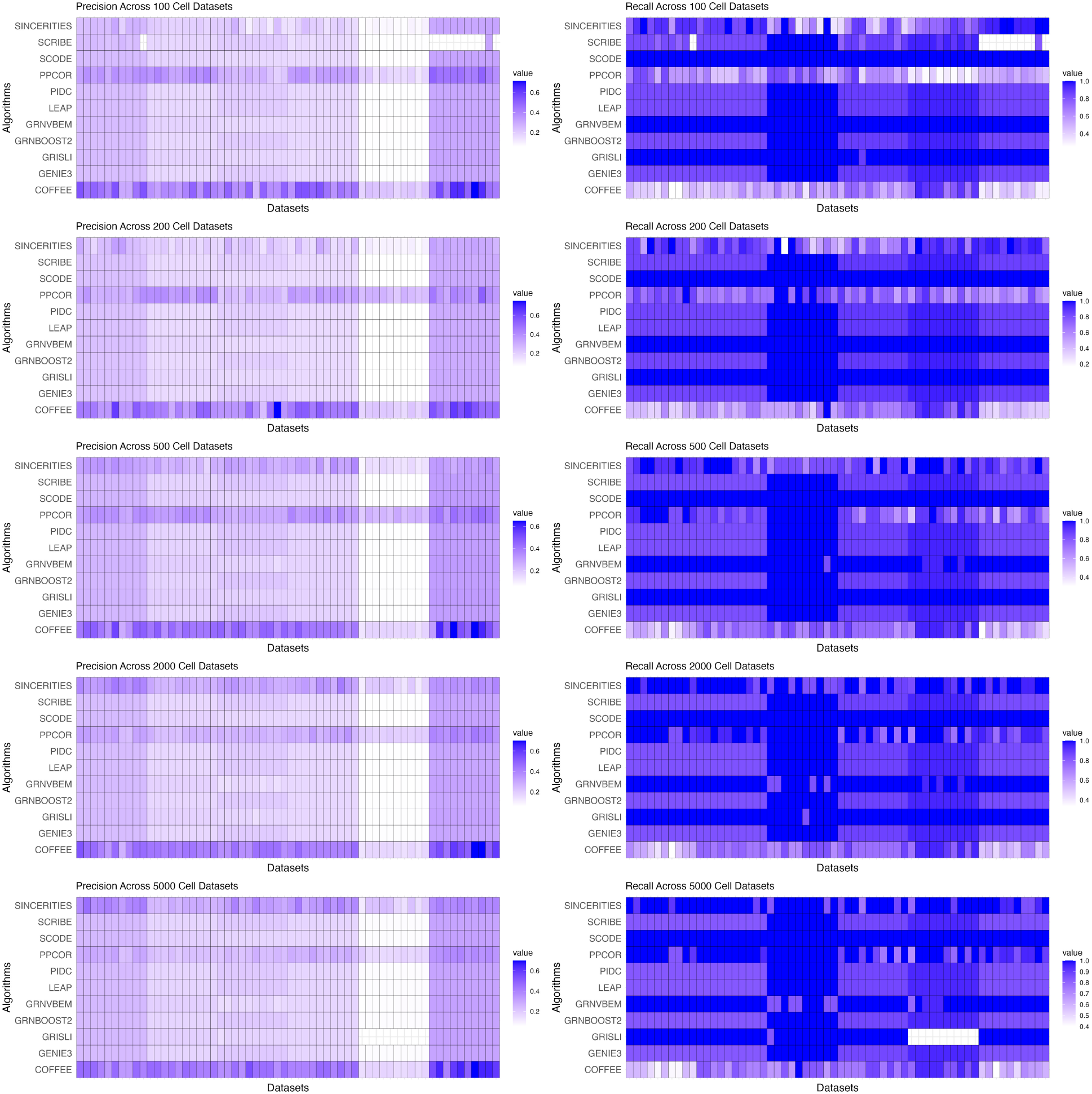
Precision & Recall performance on Synthetic datasets grouped By Size.

### 4.2 Curated Datasets

We next evaluated the performance of COFFEE on the Curated datasets from the BEELINE evaluation framework. Each dataset contained 2000 cells, so the threshold used for COFFEE was the previously identified optimal one of 0.65. We evaluated COFFEE’s performance using precision, recall and F-Score.

The precision-recall analysis for Curated datasets can be seen in Figure 4. Similar to the Synthetic sets, COFFEE showcases higher precision compared to the baseline algorithms in three of the four datasets. This demonstrates COFFEE’s stability across differing datasets; even with Curated data, the consensus approach is able to capture high confidence edges when compared to the ground truth data.

**Figure 4:**
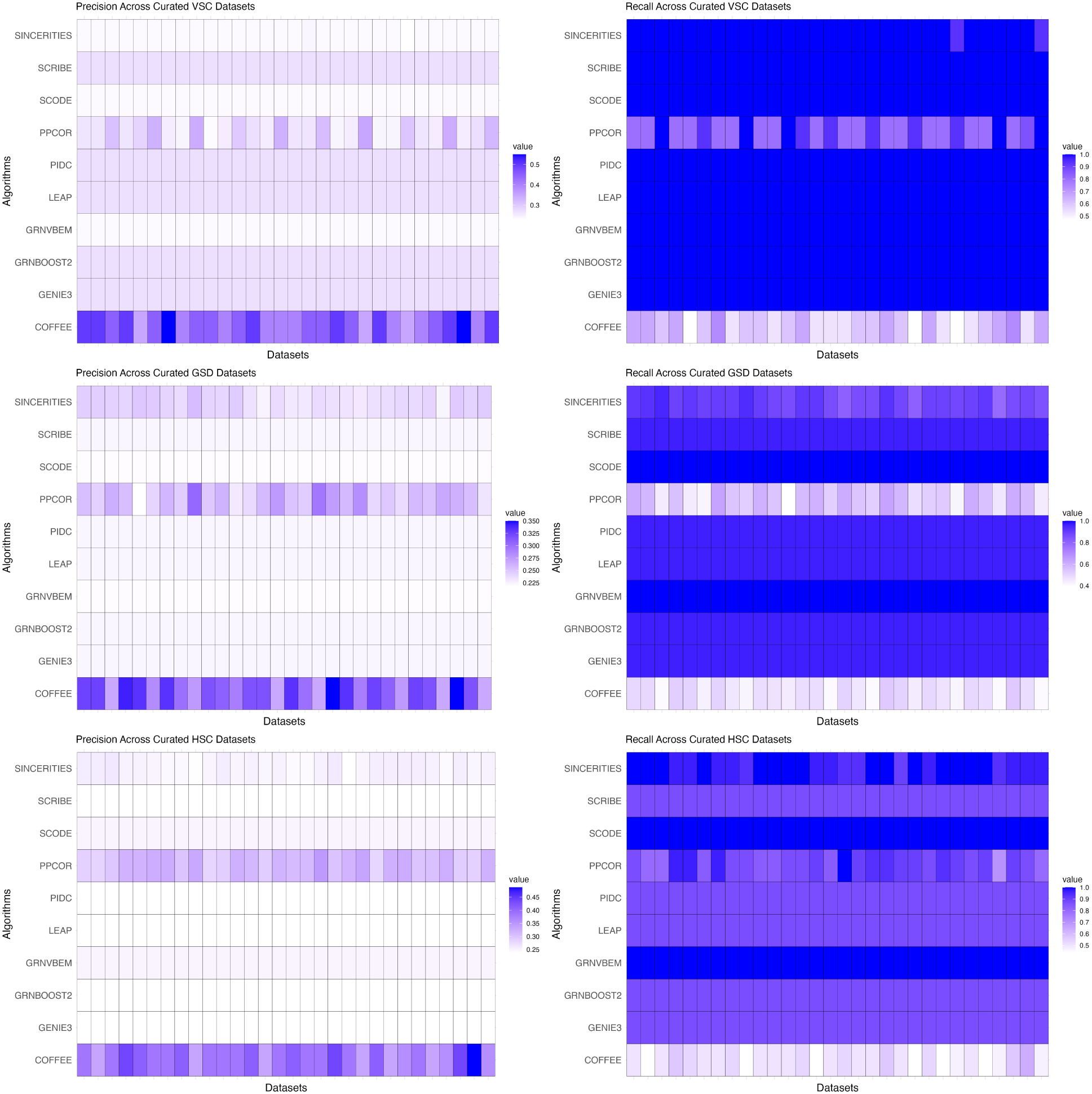
Precision & Recall Performance on Curated Datasets.

We further explored COFFEE’s performance by F-Score against the baseline algorithms. These results are visualized in Figure 5. We can observe that COFFEE performs very well compared to the baseline algorithms in terms of F-Score in three of the four datasets. Much like the Synthetic datasets, we see high variation in the baseline algorithms performance, even when applied to Curated sets. For example, SCODE performed better than most other algorithms in the GSD Curated dataset, but performed the worst in the VSC set. COFFEE was able to perform better than most other algorithms in three of the four datasets.

**Figure 5:**
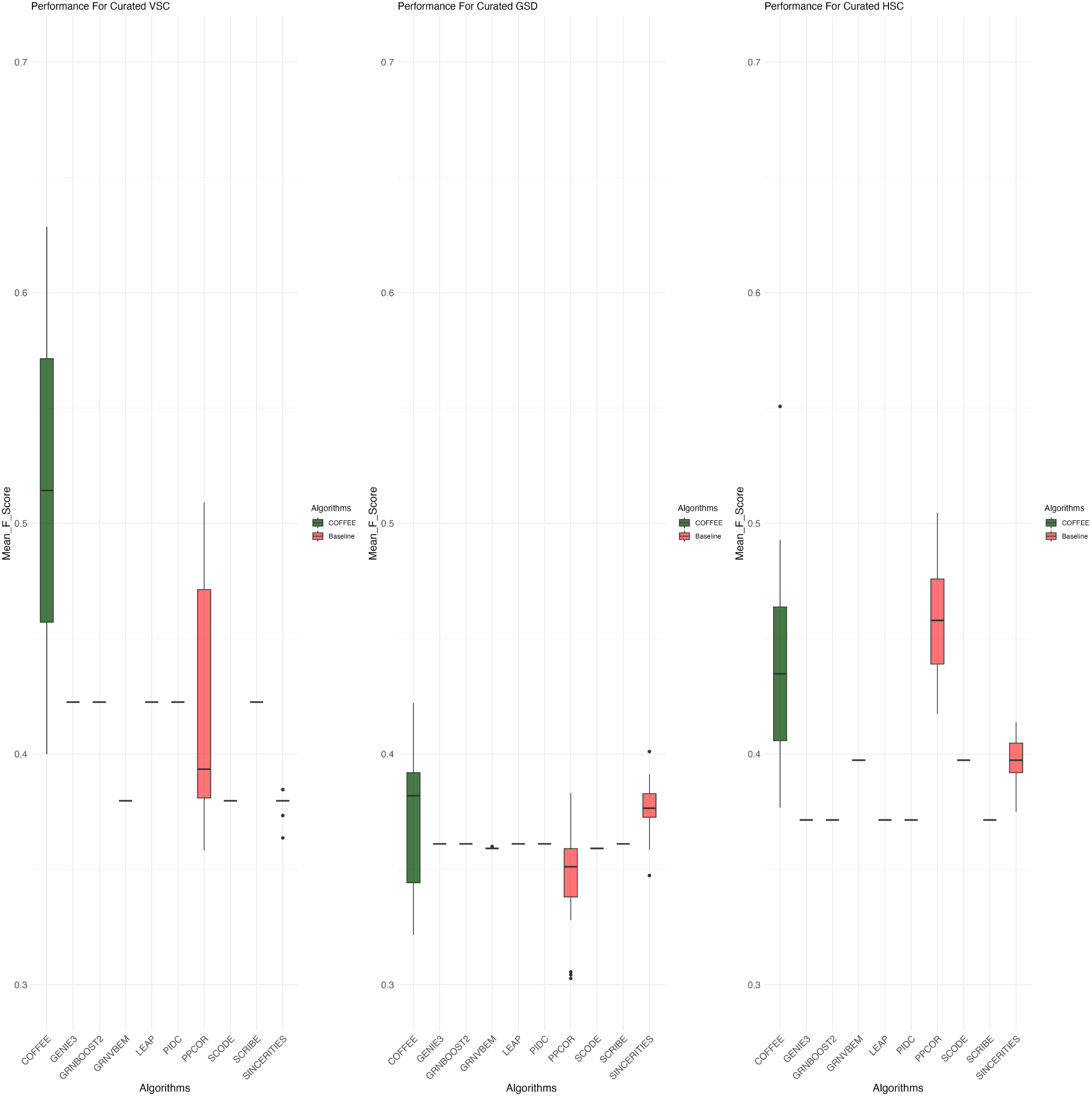
F-Score performance on Curated Datasets.

### 4.3 Experimental Datasets

Furthermore, we evaluated the performance of COFFEE on four of the Experimental datasets from the BEELINE evaluation framework. Each dataset contained a variable number of cells, and so we used a threshold of 0.65. We evaluated COFFEE’s performance using precision, recall and F-Score.

To be consistent with BEELINE’s evaluation framework, we evaluated the Experimental datasets on cell-type specific and non-specific networks. All datasets were collected from Chip-Seq protocol, as outlined in BEELINE [2].

Figures 6 and 7 display the results of COFFEE compared to the baseline algorithms. We noticed that some algorithms did not predict any edges for certain datasets, while they did predict some edges for the others. Therefore, each COFFEE run contained a differing number of algorithms across datasets.

**Figure 6:**
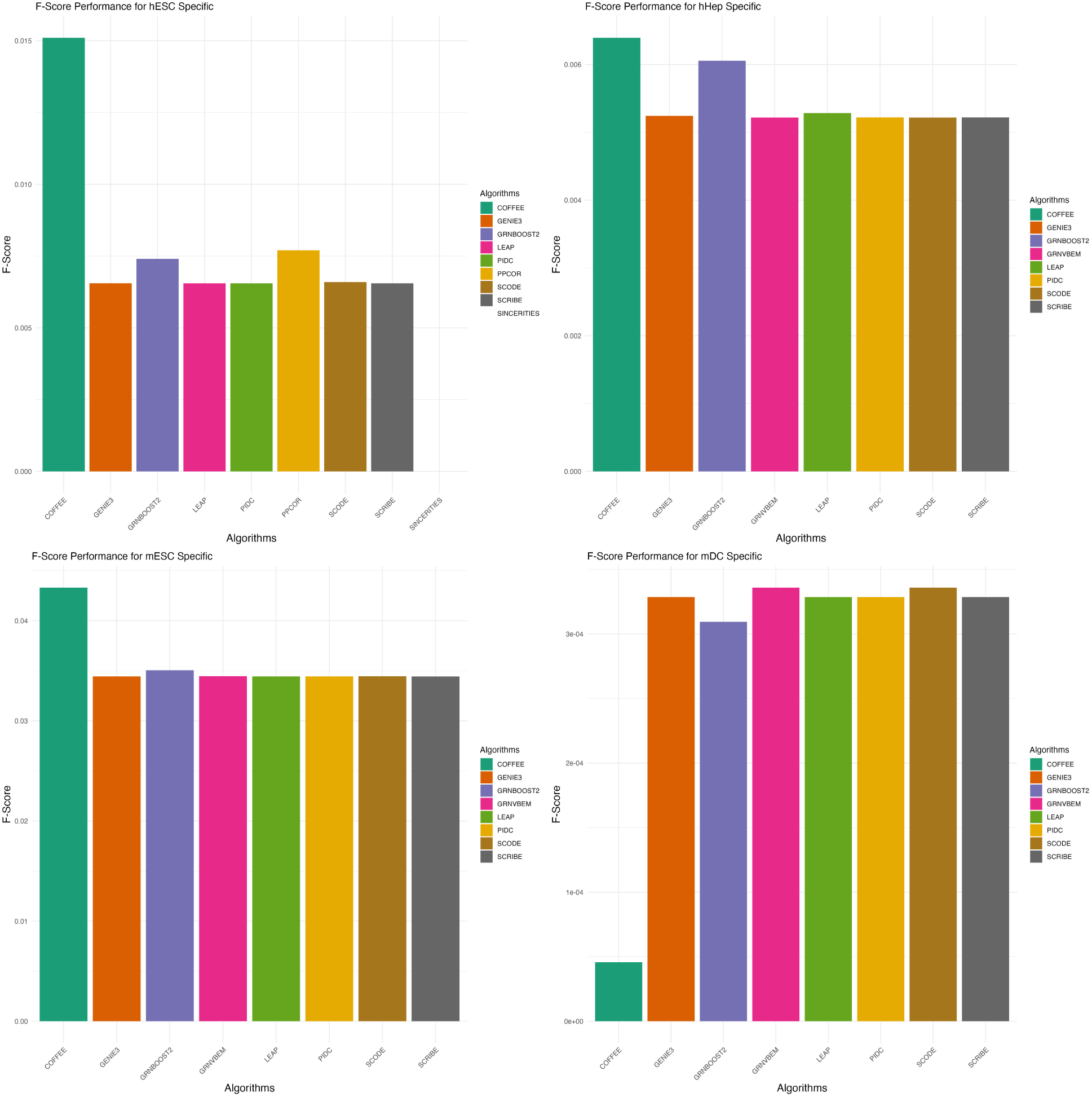
F-Score performance on Experimental datasets using Cell-Type Specific Ground Truth.

**Figure 7:**
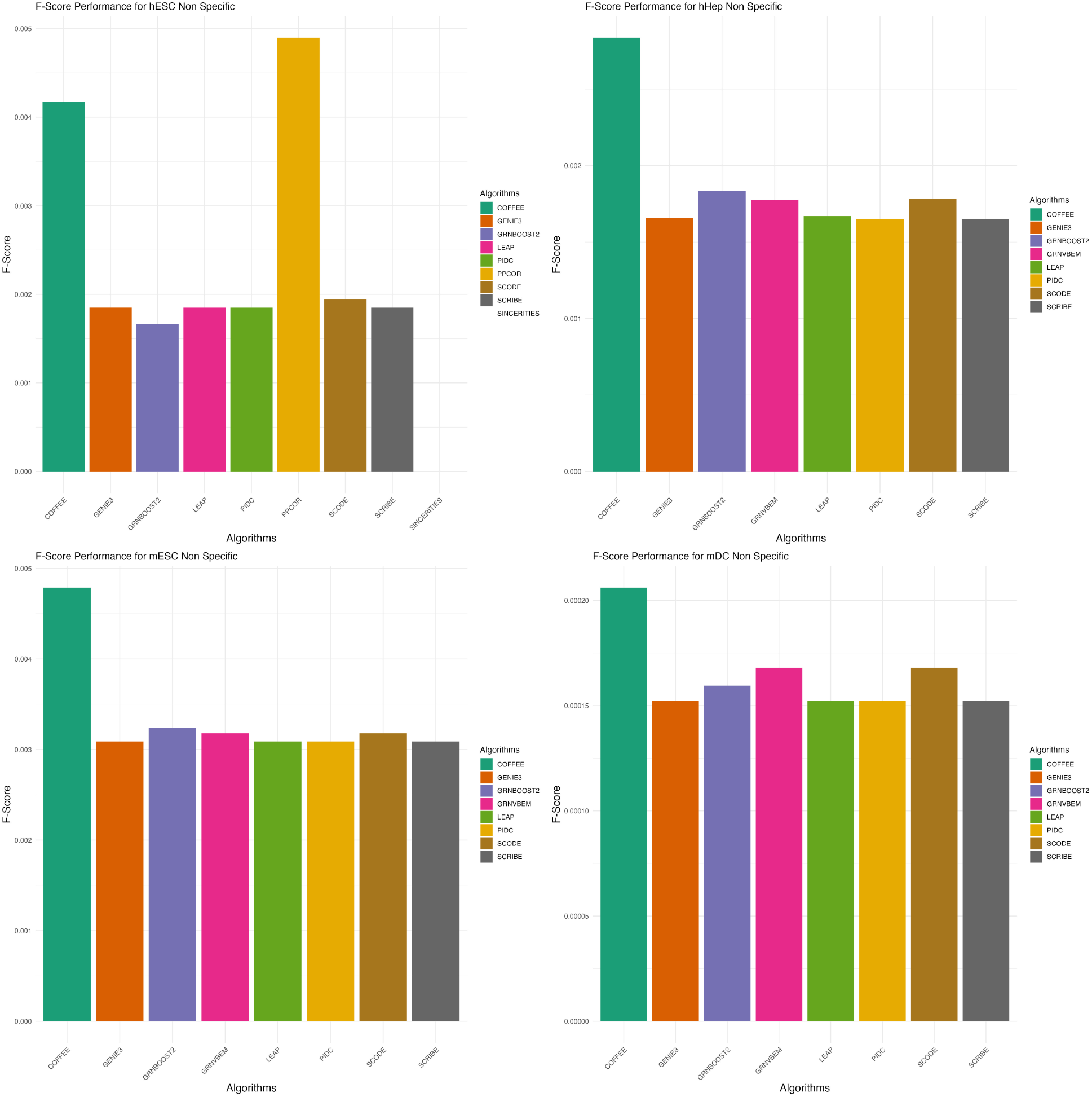
F-Score performance on Experimental datasets using non-Cell-Type Specific Ground Truth.

Across the majority of datasets and both the cell-type specific ground truth and non-specific ground truths, we see that COFFEE performs better than the baseline algorithms in terms of F-Score. However, for the mDC dataset, COFFEE performed poorly; the BEELINE evaluation framework also noted that mDC was an outlier in their performance evaluations, with several algorithms demonstrating worse than normal performance [2]. This could be due to mDC having a much higher density in their gold-standard network, making it difficult for algorithms to identify high confidence edges.

## 5 Cell-Type specific inference

Existing GRN inference methods for scRNA-seq data have not been specifically designed for cell-type specific datasets and networks. These methods do not consider the global regulatory structure of cell-types within tissue, and therefore may not be representative of key regulatory interactions [17]. To test the consensus method for cell-type specific inference, we first tried a two-level consensus approach.

On each of the Synthetic datasets used to evaluate the performance of COFFEE, we partitioned the dataset into cell types using Seurat clustering, and filtered the gene expression matrices and pseudotime data accordingly [37]. From here, we ran COFFEE with all 10 baseline algorithms; once the networks were obtained for each cell type, we ran the Borda point algorithm again, this time integrating edges from each cell type to have one composite network for the dataset. We then compared this second level consensus network to the COFFEE algorithm ran on the whole Synthetic datasets without cell-type partitioning.

We see in figure 8 that using the second level consensus approach drastically decreases the performance of COFFEE. We conclude from this analysis that the networks from individual cell-types are not entirely representative of the true global regulatory structure; therefore, a different approach is required for cell-type specific inference, that takes into account cell-lineage information [5].

**Figure 8:**
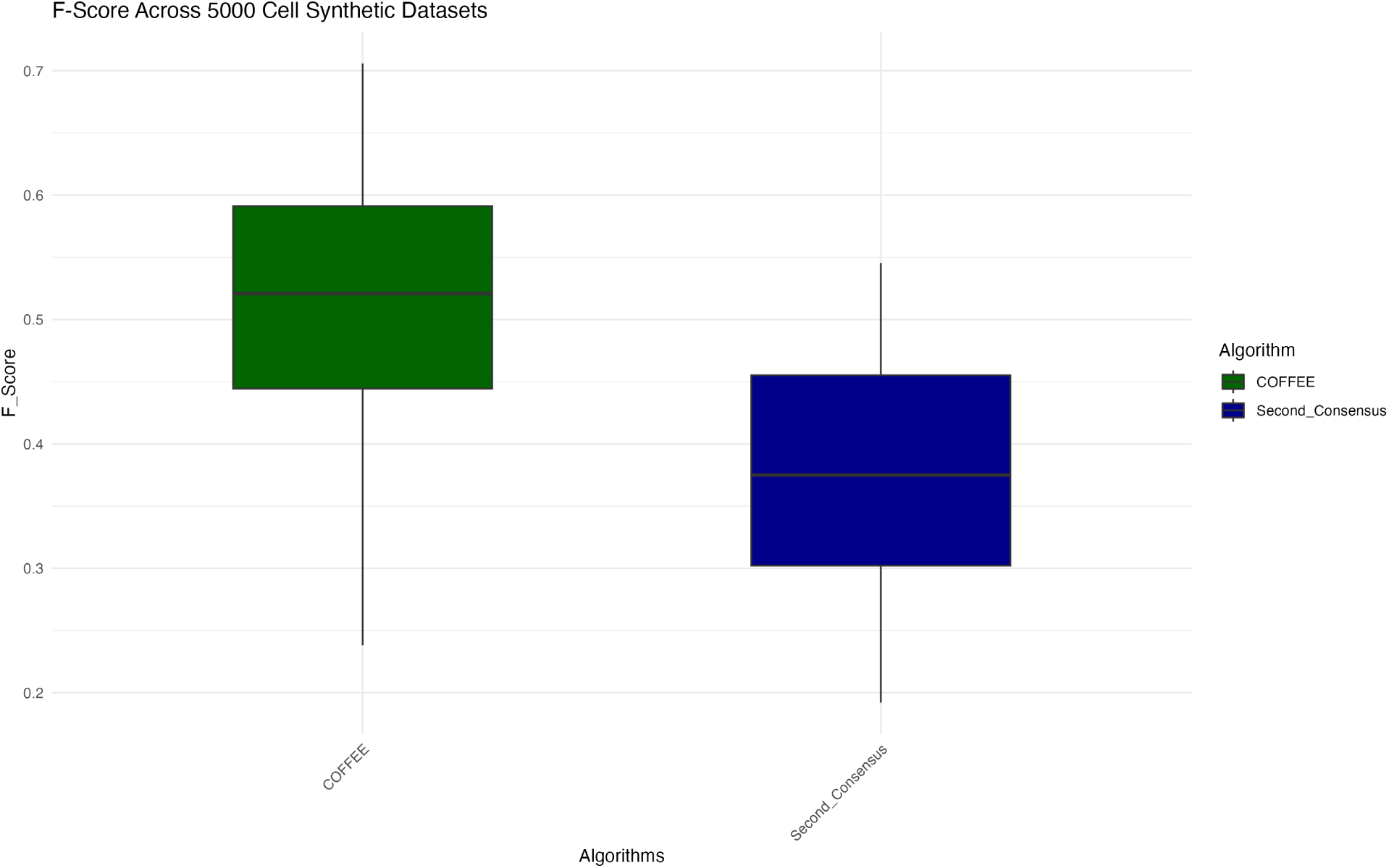
F-Score performance for Second Level Consensus vs COFFEE on Synthetic datasets.

To analyze the performance of COFFEE against scMTNI for cell-type specific performance, we used a dataset that scMTNI was initially benchmarked on [5]. The dataset is from a Human Fetal Hematopoietic cell study that studied the regulatory dynamics during human development for multiple human blood cell types [26]. In concordance with scMTNI’s benchmarking process, we used the annotated lineages clusters, which were: Hematopoietic Stem Cell and Multi-Potent Progenitor (HSC-MPP), MKerythroid-mast progenitors combined with cycling MEMPs (MEMPs), granulocytic progenitors (GPs), and lymphoid-myeloid progenitors (LMPs) [5]..

We used the single cell expression matrix provided by the authors of scMTNI to run the four algorithms in COFFEE. Similar to the BEELINE framework, we calculated the 500 most varying genes across pseudotime points using Slingshot per cell type. [9], using one lineage. These 500 genes were used to infer the GRNs for COFFEE.

For evaluation, we used the Cus-KO gold standard network described in scMTNI [5]. We chose this dataset to be gold standard since it was the network that scMTNI had the best performance metrics with [5]. Cus-KO contains interactions from the knock-down based GM12878 lymphoblastoid cell line downloaded from Cusanovich et al [38]. We filtered the gold-standard network to only contain interactions with a pvalue < 0.01. Additionally, we filtered the inferred networks from both scMTNI and COFFEE to contain only genes present in the gold standard network. Finally, we selected the top 1000 edges from the scMTNI inferred networks, and performed a sensitivity analysis for the best COFFEE threshold.

In Figure 9, we see the performance of COFFEE for various thresholds when compared to scMTNI. In two cell types, LMP and MEMP, COFFEE showcases a better performance by F-Score. However, scMTNI performes significantly better on the GPs and HSC-MPP cell types. We also noted that there is little to no effect with the COFFEE threshold on the performance of the algorithm.

**Figure 9:**
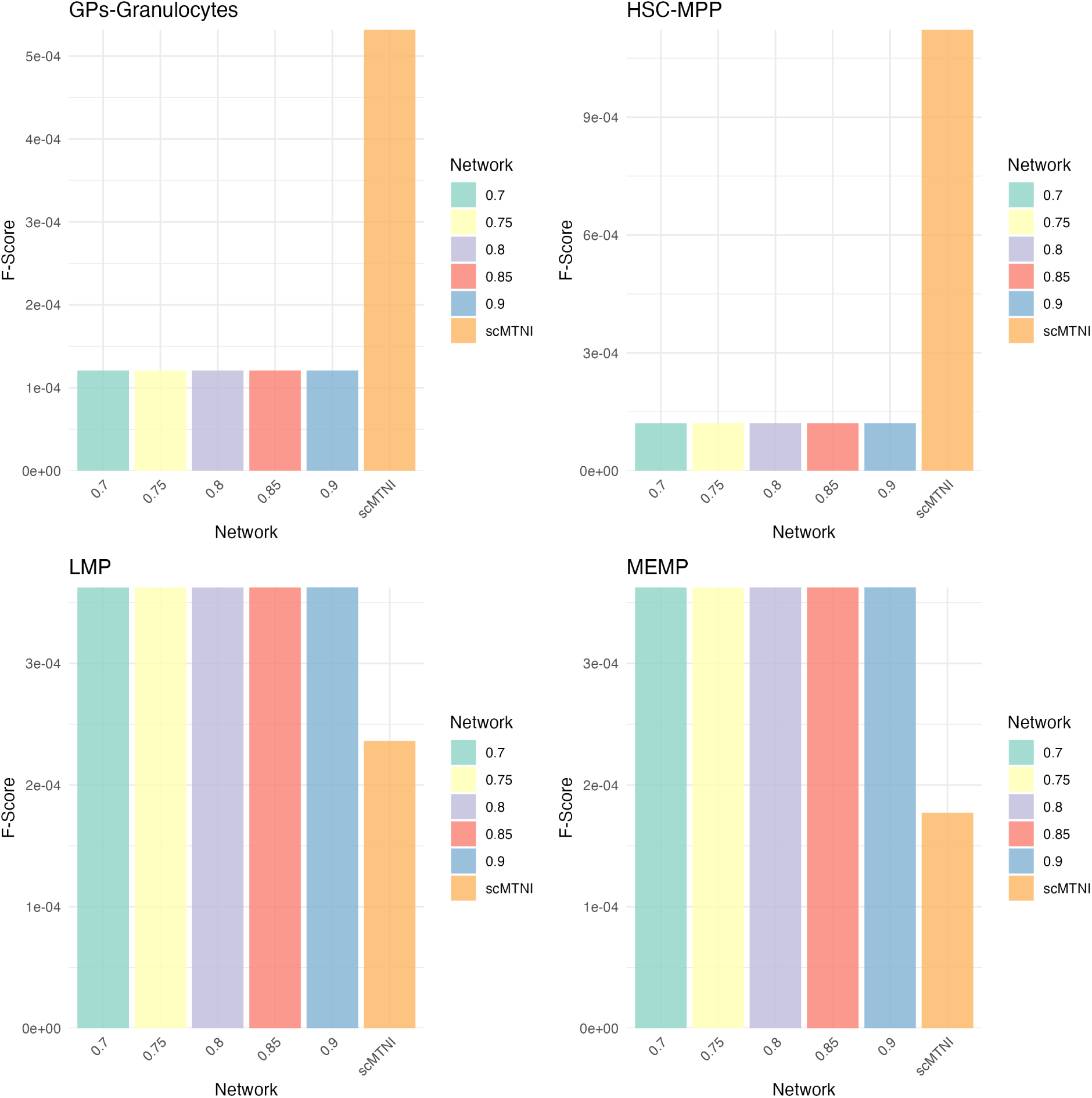
F-Score for each Cell Type across COFFEE thresholds and scMTNI.

With the cell-type specific prioritized algorithm, which we call COFFEE+scMTNI, we see that a consensus approach is able to improve the performance of scMTNI on all cell types. Thus, we can determine that the individual four algorithms predict true interactions that were not initially learned by scMTNI. We evaluated COFFEE+scMTNI on the Human Fetal Hematopoietic as described earlier [26]. The results of this analysis are shown in Figure 10.

**Figure 10:**
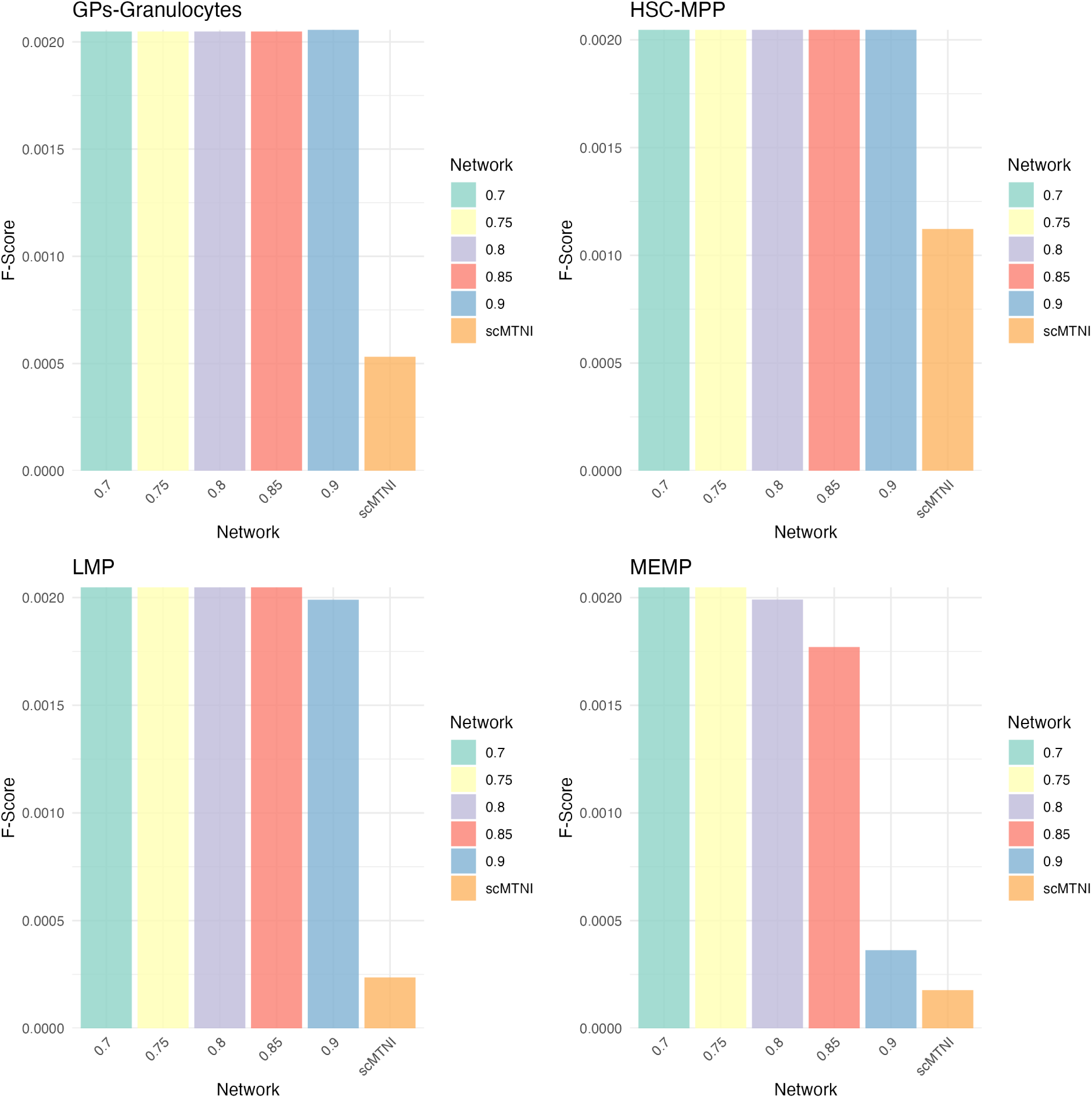
F-Score for each Cell-Type across COFFEE thresholds and COFFEE+scMTNI.

## 6 Discussion

From our analysis, we are able to conclude that a consensus approach for single cell GRN inference has several advantages. Across various types of datasets, COFFEE demonstrated improved performance in determining true interactions when compared to gold standard datasets.

For Synthetic datasets, we determined the ideal threshold to select edges from the consensus network generated by COFFEE. We found that for larger datasets, including more edges improved the performance of the algorithm, while for smaller datasets, a higher threshold maximized the F-Score when compared to the gold standard. However, users may want to adjust the threshold value based on the statistic they want to maximize; increasing the consensus threshold will lead to a higher precision, while decreasing the threshold will lead to a higher recall. The suggested thresholds in Table 6 are based on the F-Score. Users ideally should run COFFEE with various threshold values and evaluate the performance on their particular dataset.

We also found that there was significant variation between algorithm performance on dataset size. For example, SINCERITIES did not perform as well as the other baseline algorithms on smaller datasets, but performed extremely well on larger datasets in terms of F-Score. The primary advantage of COFFEE is that the algorithm has consistent high performance across multiple types of datasets; this is inherent in the algorithm design, as edges appearing in multiple GRN inferred networks will have higher confidence scores in the final consensus network. This approach establishes that the consensus approach will work across several different kinds of datasets, particularly with a large selection of algorithms to integrate.

This effect is even more pronounced when analyzing Curated and Experimental datasets. Despite the number of cells being the same across Curated datasets, different algorithms performed better on some datasets than others. This establishes that users cannot go by dataset size alone when choosing an optimal algorithm. Additionally, algorithms that require pseudotime data as input are more likely to suffer in their performance if the pseudotime data is inaccurate, or incorrectly computed [2]. While Pratapa et al. in their benchmarking study make suggestions for which algorithms to use for various datasets, a far safer approach would be to use a consensus based method. We see that across Curated datasets, COFFEE has high F-Score values. In contrast, we see that each Curated dataset had a different top performer. Without considering COFFEE, the top performer for VSC was GENIE3, PPCOR for HSC, and SINCERITIES for GSD.

The variation in algorithm performance across datasets makes a consensus method such as COFFEE a safe choice to consistently infer high quality networks.

On the Experimental Datasets, we chose to evaluate on Cell-Type specific and Non-Specific ground truths, as was done in the BEELINE evaluation framework [2]. From this analysis, we noticed that while the baseline algorithms performed poorly on the Non-Specific networks, COFFEE performed significantly better. However for the Cell-Type Specific ground truth data, COFFEE performed better generally, but there was less difference between the baseline and consensus methods in terms of F-Score. This suggests that COFFEE without modifications is not as well equipped to infer more specific interactions, rather than general interactions collected by organism level Chip-Seq networks. This trend is also seen when evaluating COFFEE based on Early Precision Ratio, or EPR. EPR is defined as the precision of the top *k* edges when compared to the ground truth, where *k* is the number of interactions present in the filtered ground truth. The reasoning behind this was that experimental groups would primarily be interested in only high confidence edges from a network [2]. COFFEE demonstrates a higher EPR, and a much better EPR for the Non-Specific networks when compared to the Cell-Type Specific networks. We show this result in supplementary figures **??** and **??**. This finding motivated us to adapt the COFFEE algorithm for cell-type specific inference, by developing a prioritized consensus method.

We see that a modified COFFEE algorithm is able to improve the performance of a well established cell-type specific GRN inference algorithm. Despite the lack of the additional information that scMTNI takes as input, COFFEE with four algorithms had improved performance on two out of four cell-types on the benchmarking dataset. We were also able to substantially improve the performance of scMTNI by incorporating a prioritized consensus approach, where interactions predicted by scMTNI were all initialized with a raw count of 1. We anticipate that the modified consensus approach has the potential to be very effective for cell-type specific GRN inference as more methods are made available to the community.

We conclude that while a regular consensus approach for bulk RNA-seq data has performed well in prior studies, a modified consensus approach is warranted for scRNA-seq data. The best practice method for bulk RNA-seq GRN inference is to incorporate as many algorithms as possible, as this demonstrates improved performance [14]. However, we see that a consensus approach for scRNA-seq data requires more careful selection of the algorithms incorporated for integration. In supplementary file **??**, we demonstrate that across Synthetic datasets of differing sizes, distinct algorithm combinations leads to variable performance. From this result we establish that simply including every GRN algorithm available will not necessarily lead to the best performance in the consensus approach. Therefore, the user needs to exercise discretion in choosing the best algorithm combination for their individual needs, and more benchmarking in this area needs to be done in order to make specific recommendations.

The primary limitation of a consensus based approach is that it is only as strong as its underlying algorithms. As more improved GRN inference algorithms emerge, it will be difficult to determine what will be the best algorithm for any given dataset. With this point in mind, we continue to see a consensus based approach to be beneficial for the community when there is uncertainty in choosing the best algorithms to use.

## 7 Conclusion

In this paper, we present COFFEE, a Borda voting based consensus algorithm for GRN inference on scRNA-seq data. COFFEE has demonstrated improved and consistent performance across Synthetic, Curated and Experimental datasets. Additionally, a modified consensus based approach for cell-type specific GRN inference has shown a promising ability to improve performance on existing state-of-the-art methods, by augmenting important gene interactions. As future work improves the landscape of cell-type specific GRN inference, this will necessitate a weighted consensus algorithm approach to merge the predictions of the sets of cell-type specific and non-cell-type specific algorithms for robust GRN inference.

## 8 Data Availability

To perform the evaluation for COFFEE against the 10 baseline algorithms, we used data from the BEELINE evaluation framework [2]. The BEELINE data may be downloaded here: https://zenodo.org/records/3378976

To perform COFFEE experiments for the cell-type specific datasets, we used the data from scMTNI’s evaluation [5]. The scMTNI data may be downloaded here: https://zenodo.org/records/7879228

