## Supplementary Figures for "COFFEE: Consensus Single Cell-Type Specific Inference for Gene Regulatory Networks"

A PREPRINT

**Musaddiq K Lodi***
Integrative Life Sciences
Virginia Commonwealth University
Richmond, VA 23284


**Anna Chernikov**
Center for Biological Data Science
Virginia Commonwealth University
Richmond, VA 23284


**Preetam Ghosh**
Department of Computer Science
Virginia Commonwealth University
Richmond, VA 23284


### 1 Supplementary Figures

*

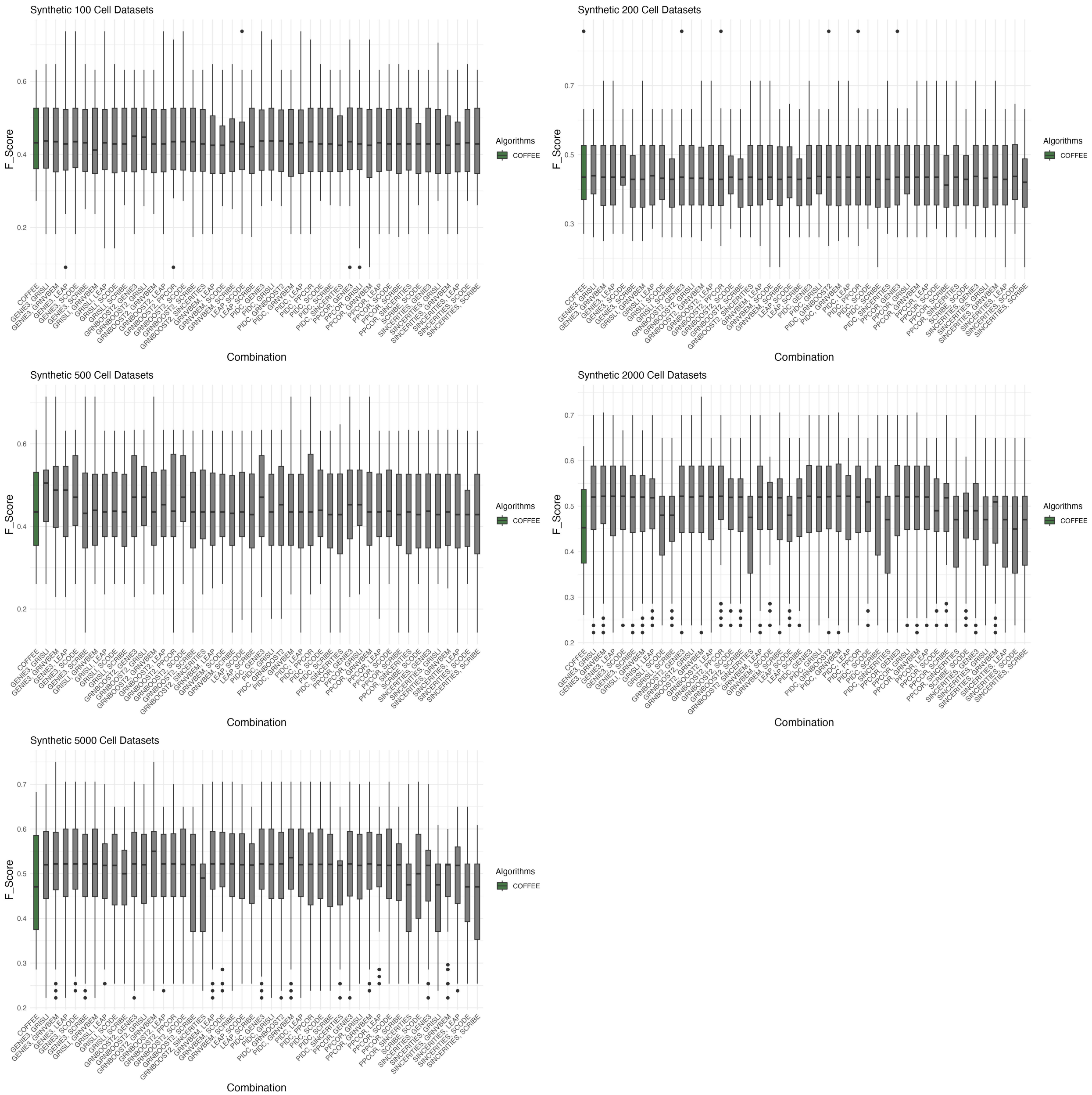


Figure S1: F-Score Performance for Different Combinations of Algorithms in COFFEE vs COFFEE with all 10 Algorithms, in green. The labels correspond to algorithms removed from the original 10 in the COFFEE framework

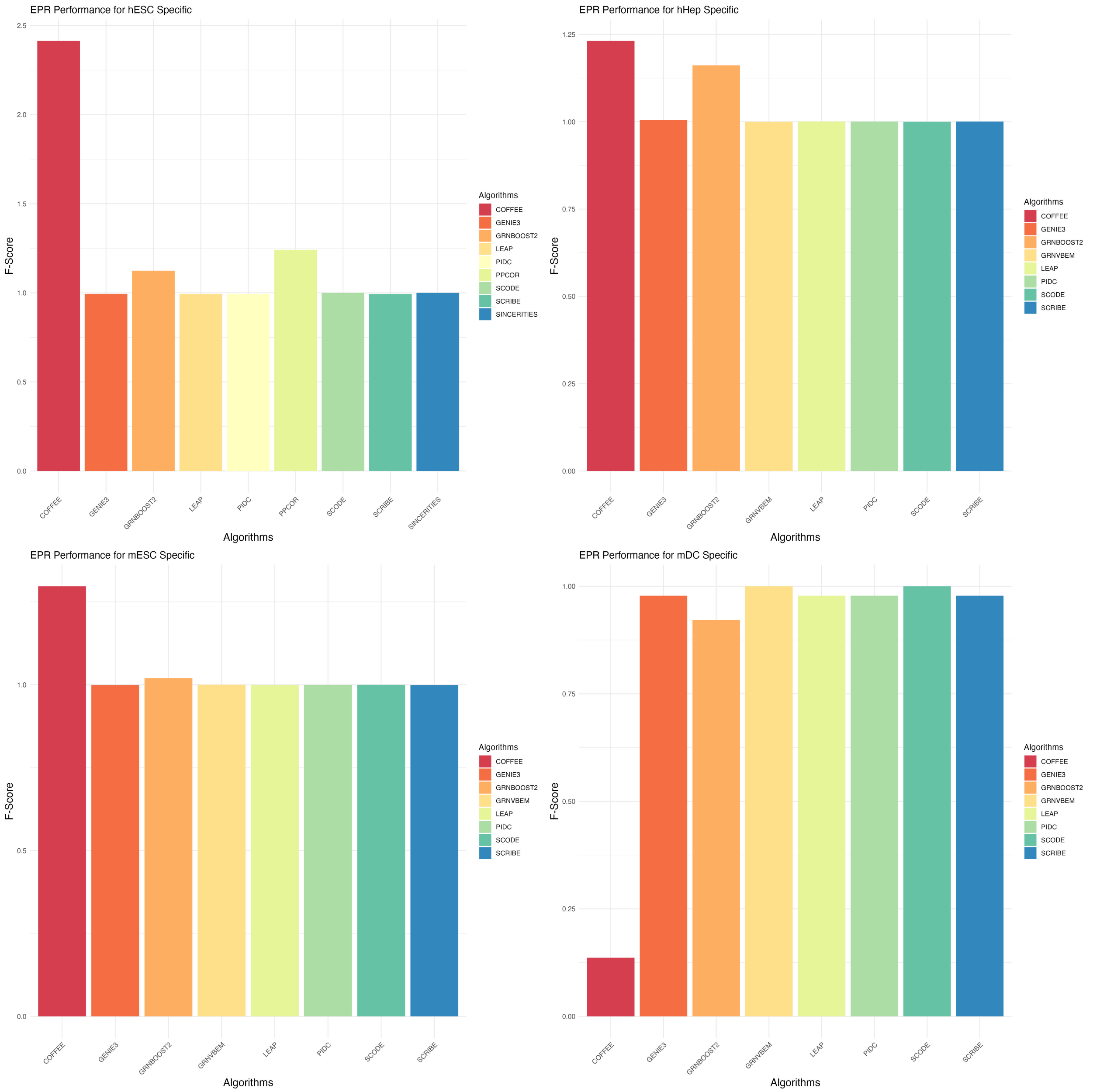


Figure S2: EPR Performance Across Experimental Datasets, Cell-Type Specific Ground Truth

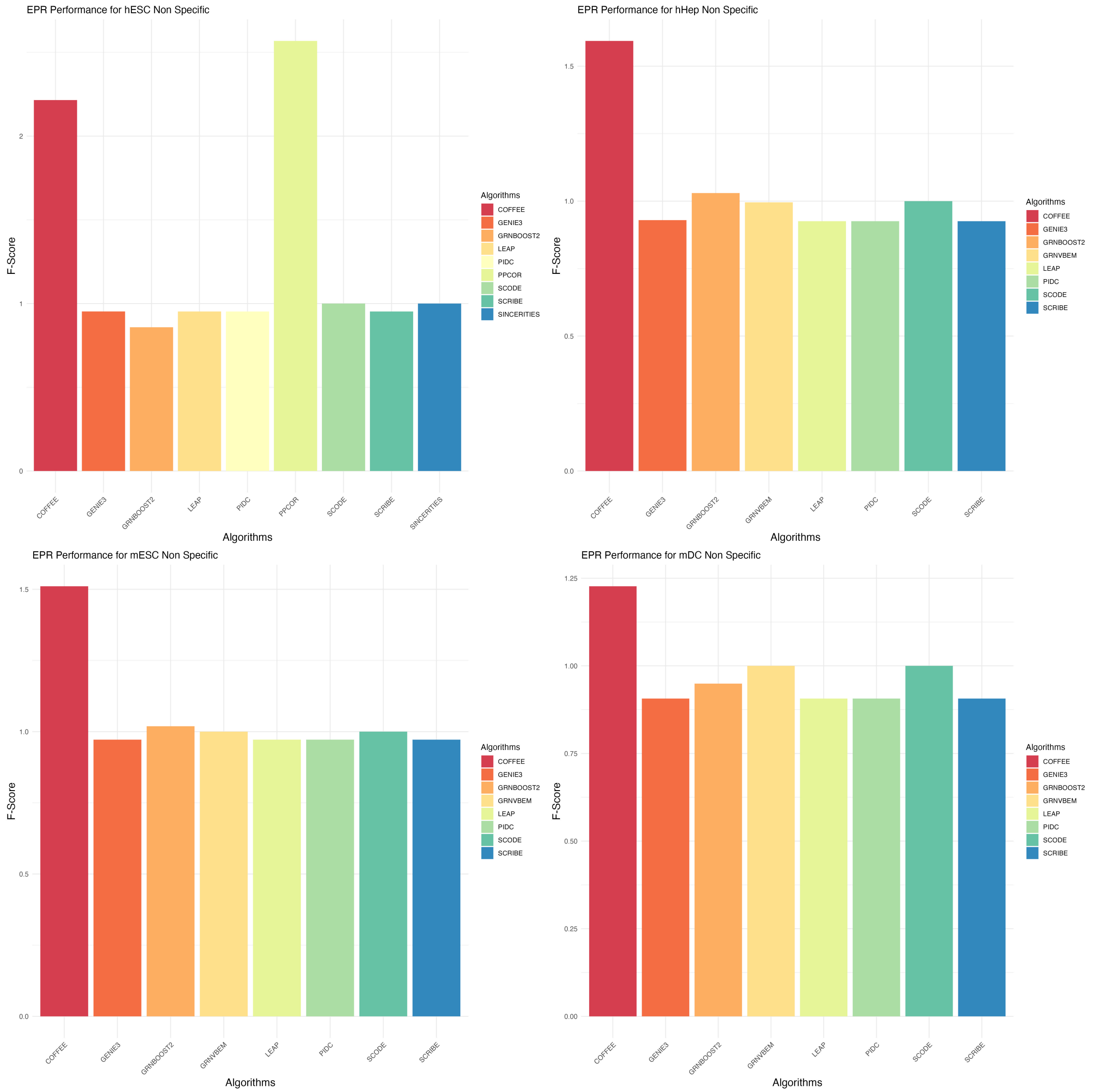


Figure S3: EPR Performance Across Experimental Datasets, Non-Specific Ground Truth
